# Acute appendicitis manifests as two microbiome state types with oral pathogens influencing severity

**DOI:** 10.1101/2022.04.13.488268

**Authors:** Marcus Blohs, Alexander Mahnert, Kevin Brunnader, Christina Flucher, Christoph Castellani, Holger Till, Georg Singer, Christine Moissl-Eichinger

## Abstract

Mounting evidence suggests that acute appendicitis (AA) is not one but two diseases: complicated appendicitis, which is associated with necrosis leading to perforation or periappendicular abscess, and uncomplicated appendicitis, which does not necessarily result in perforation. Even though AA is the most frequent cause of surgery from abdominal pain, little is known about the origins and etiopathogenesis of this disease, much less regarding the different disease types.

In this study, we investigated the microbiome of samples from the appendix, rectum and peritoneum of 60 children and adolescents with AA to assess the composition and potential function of bacteria, archaea and fungi. The analysis of the appendix microbial community revealed a shift depending on the severity of the AA. This shift was reflected by two major community state types that represented the complicated and uncomplicated cases. We could demonstrate that complicated, but not uncomplicated, appendicitis is associated with a significant local expansion of oral, bacterial pathogens in the appendix, most strongly influenced by necrotizing *Fusobacterium* spp., *Porphyromonas* and *Parvimonas*. Uncomplicated appendicitis, however, was characterised by gut-associated microbiomes. Our findings support the hypothesis that two disease types exist in AA, which cannot be distinguished beyond doubt using standard clinical characterization methods or by analysis of the patient’s rectal microbiome. An advanced microbiome diagnosis, however, could improve non-surgical treatment of uncomplicated AA.

**Importance:** With a lifetime risk of up to 17%, acute appendicitis is one of the most frequent causes of emergency abdominal surgery in westernized countries. Latest literature reports suggests that appendicitis manifests in two disease types: complicated and uncomplicated appendicitis with different, yet unknown, etiopathogenesis.

In this study, we investigated the microbial composition (bacteria, archaea and fungi) from 60 children and adolescents that were diagnosed with acute appendicitis. Appendix, rectal and peritoneal samples were analysed using amplicon and metagenomic sequencing. Our results suggest that acute appendicitis manifests in three microbial state types that reflect complicated and uncomplicated appendicitis as well as special cases that are caused by bacterial overgrowth. Strikingly, uncomplicated appendicitis appears to be caused by gut-associated pathogens while complicated appendicitis is driven by oral-associated microbes such as Fusobacterium sp. or Porphyromonas sp. The findings provided in our study are of special interest to understand the etiopathogenesis of both complicated and uncomplicated appendicitis.

## Introduction

With approximately 100 cases per 100,000 person-years, acute appendicitis (AA) is the most common reason for emergency abdominal surgery in westernised countries. The lifetime risk of developing AA is estimated at between 6–17%, depending on a person’s sex, life expectancy, region and socioeconomic status (1–4). While a distinct morphological succession can be observed in the appendix during AA (Fig. 1), the etiopathogenesis of this disease is still not fully understood. Historically, appendicitis was thought to result from (temporal) luminal obstruction, followed by distention, bacterial overgrowth and increased intraluminal pressure, eventually resulting in the disintegration of the vermiform appendix wall and thereby to gangrene or perforation (5). However, this hypothesis has only limited support, as both obstruction by a faecalith and increased luminal pressure are only found in about 20% and 25% of appendicitis patients, respectively (6, 7). Furthermore, researchers have argued that AA does not necessarily result in gangrene or perforation. Appendicitis may, in fact, represent two different diseases: uncomplicated and complicated appendicitis, each with a distinct epidemiology and pathophysiology (8–10). Complicated appendicitis has been described as perforated appendicitis, periappendicular abscess, or peritonitis, which is defined as an acute inflammation of the peritoneum that occurs in addition to the infection of the appendix (11)

**FIGURE 1.**
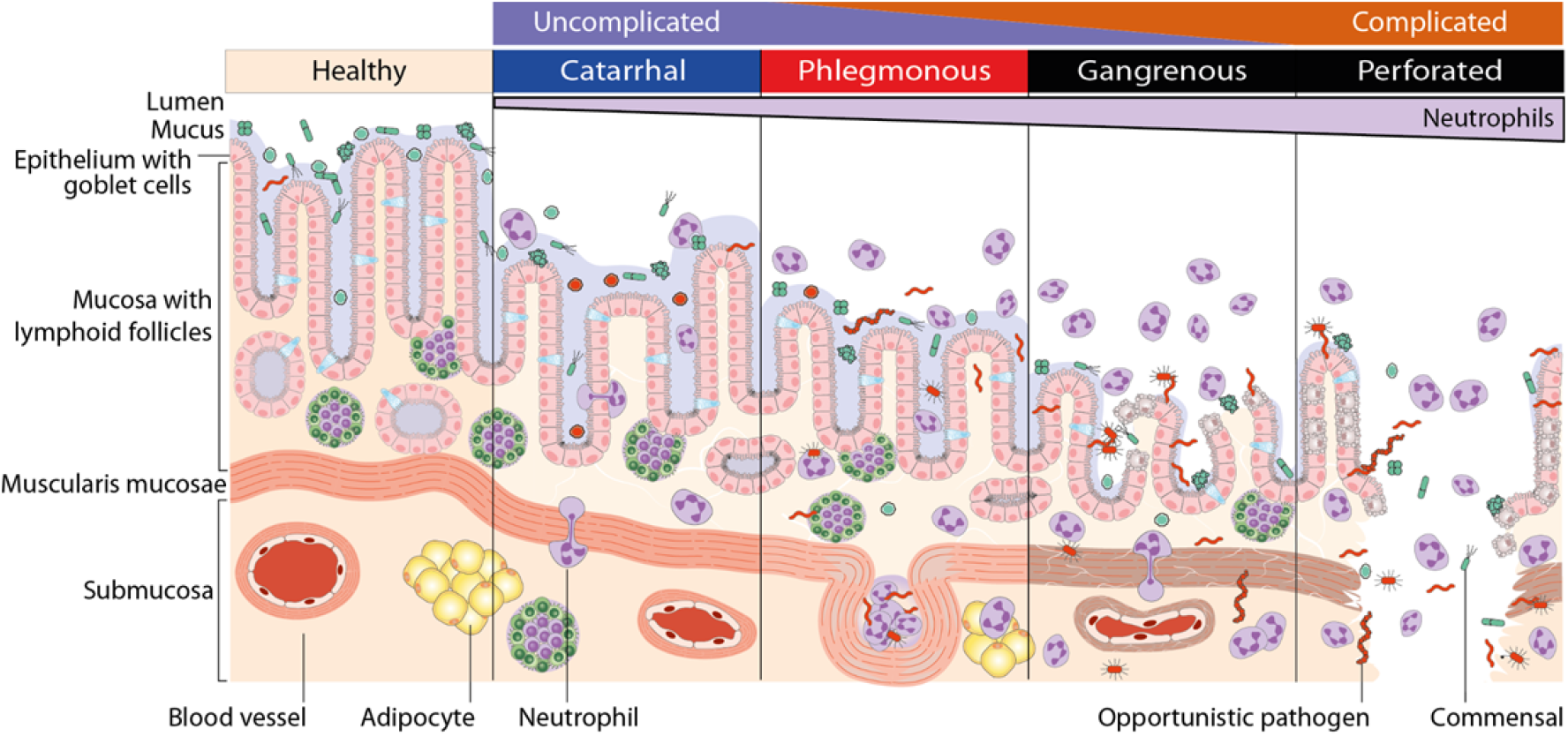
Disease stages of acute appendicitis. **Catarrhal** or early appendicitis is characterised by neutrophilic infiltration of the appendix wall, whereby neutrophils transmigrate into the lumen, caused by luminal obstruction or bacterial infection. Fluid accumulation and increased intraluminal pressure may lead to tissue distention, mucosal ulceration and bacterial passage through the epithelium. In **phlegmonous** appendicitis, the inflammation involves the entire appendix wall and leads to extensive ulceration, vascular thrombosis and frequently to intramural abscess formation. Increasing intraluminal pressure and thrombosis support bacterial tissue penetration and lead to **gangrenous** appendicitis with large areas of tissue necrosis. Terminally persisting tissue damage can result in **perforation**. (26)

Recently, mounting evidence suggests that microbial dysbiosis (12–15) together with an uncontrolled inflammatory response (16–18) drive the development of the disease. This claim is supported by the successful treatment of uncomplicated appendicitis with broad-spectrum antibiotics as an alternative to surgery (19). However, as in other gastrointestinal diseases, it is difficult to pinpoint the specific microorganisms that are responsible for the development of appendicitis. One problem is that each “healthy” microbiome is unique and may already include opportunistic pathogens that cause diseases. Another problem is the non-accessibility of healthy appendix microbiomes, as incidental appendectomies are not performed routinely.

Previous work using both culture-dependent and -independent methods suggested the existence of a tight connection between the occurrence and abundance of specific microbial taxa and AA. The latest studies based on amplicon sequencing and real-time quantitative polymerase chain reaction (RT-qPCR) identified a local expansion of oral cavity-associated microbes, including *Fusobacterium, Peptostreptococcus, Porphyromonas* and *Gemella* (13, 15, 20, 21). Other studies have connected the inflammation to food-borne pathogens such as *Campylobacter jejuni* (22). Blood *et al*. (21) even have shown that some of the aforementioned oral pathogens can potentially survive the passage through the stomach, suggesting that AA may be an infectious disease. Out of these oral microorganisms, only *Fusobacterium* spp. are recurring taxa commonly found in AA (12, 13, 15, 20, 23, 24), with *F. nucleatum* being capable of infiltrating the appendix lumen depending on the disease severity (23). In contrast to their reportedly high impact in AA, Fusobacteria were only found in 62% of the appendicitis patients described by Swidsinsky *et al*. (23) and, likewise, other pathogens are also inconsistently found among AA patients, suggesting a different/individual etiopathogenesis for AA.

Only a few studies with a limited number of patients have been carried out to investigate differences in the microbial composition at different stages of AA (20, 24, 25). As microbial causes and/or responses may be specific to the grade of disease severity, we prospectively recruited 60 AA patients, grouped them according to postoperative pathological findings and analysed the microbial composition in the appendix as well as in the peritoneum and rectum (swabs). For this purpose, we performed 16S/23S rRNA gene amplicon sequencing for all samples and shotgun metagenomic analysis of appendix samples. Below, we describe the functional and compositional relevance of bacteria, archaea and fungi with respect to different stages of AA.

## Results

### Study population

In our study, we were able to monitor microbial differences in the appendix, rectum and peritoneum throughout three severity stages of acute appendicitis (AA). Tissue samples of the vermiform appendix and rectal swabs were obtained from 60 patients diagnosed with AA (median age: 12.0, range: 3 to 17 years), while peritoneal swabs were taken from 35 of the 60 recruited participants. The final evaluation of disease severity and subgrouping of samples was performed based on a histopathological investigation as follows: subacute, catarrhal, phlegmonous, gangrenous and perforated (Tab. 1). In total, four patients without notable pathological findings (subacute samples) were excluded from further microbial analyses due to insufficient sample size. The defined patient groups were not significantly different regarding their age, sex, PAS and Alvarado score or the presence of a faecalith. However, the serum CRP concentration was significantly elevated in patients with gangrenous/perforated appendicitis as compared to those with phlegmonous appendicitis (Kruskal-Wallis test; *P* < .01). Furthermore, the number of patients treated with antibiotics was significantly higher in gangrenous/perforated appendicitis (chi-square test; *P* < .05). A total of 20 patients (33%) were administered antibiotics intravenously immediately before or during surgery, depending on the surgeon’s assessment. We did not expect these antibiotics to affect the microbiome due to the short interval between administration and sampling. Even differences in bacterial beta diversity in relation to antibiotics treatment (amplicon data: PERMANOVA - Bray-Curtis, *P* = .048; weighted UniFrac, *P* = .029) are clear confounding factors of the disease severity (antibiotics are applied significantly more often in complicated cases; Tab. 1) and, therefore, were not considered to have a substantial effect on microbial diversity.

**TABLE 1.** Clinical and demographic data for the study cohort grouped by pathological findings of acute appendicitis.

| Study population | subacute<br>(n = 4) | catarrhal<br>(n = 14) | phlegmonous<br>(n = 31) | gangrenous/<br>perforated<br>(n = 11) | P-value |
| --- | --- | --- | --- | --- | --- |
| Age (yrs)<sup>a</sup> ± SD | 10.0 ± 4.2 | 10.3 ± 3.0 | 10.9 ± 3.2 | 10.3 ± 4.7 | .63 |
| Male sex, n (%) | 2 (50) | 9 (64) | 26 (84) | 5 (45) | .12 |
| PAS score (0-10)<sup>a</sup> ± SD | 6.5 ± 1.9 | 6.6 ± 2.3 | 6.7 ± 2.3 | 8.7 ± 1.5 | .19 |
| Alvarado score (0-12)<sup>a</sup> ± SD | 6.8 ± 1.9 | 6.6 ± 2.6 | 6.6 ± 2.4 | 8.2 ± 2.7 | .20 |
| Leukocyte count (10<sup>9</sup>/L)<sup>a</sup> ± SD | 15.4 ± 8.4 | 13.2 ± 4.6 | 14.1 ± 5.6 | 14.5 ± 4.6 | .88 |
| C-reactive protein (mg/l)<sup>a</sup> ± SD | 68.5 ± 63.2 | 37.9 ± 43.2 | *23.5 ± 35.7 | 117.2 ± 101.3 | **.002** |
| Faecalith detected, n (%) | 0 (0) | 1 (7) | 5 (16) | 1 (9) | .69 |
| Antibiotic treatment, n (%) | *0 (0) | *2 (14) | *10 (32) | 8 (73) | **.007** |

<sup>a</sup>, mean values; \*, sign. different from gangrenous/perforated; SD, standard deviation

### Bacteriome in acute appendicitis patients

We first evaluated the impact of the bacterial microbiome upon disease progression for the three sample types (appendix, peritoneal and rectal samples). Based on 16S rRNA gene amplicon sequencing of the V4 region, we identified a total of 2,158 ASVs in the dataset after quality control, filtering and SRS normalisation. All three sample types were dominated by the phyla Firmicutes (see Fig. 2A; <u>A</u>ppendix: 34.4%, <u>P</u>eritoneum: 32.9%, <u>R</u>ectal: 50.0%) and Bacteroidota (A: 31.1%, P: 15.5%, R: 27.5%), while Proteobacteria were identified as the second most abundant phylum in peritoneum samples (29.7%) but were less abundant in the other two sample types (A: 12.4%, R: 6.1%). Fusobacteriota were solely represented by the genus *Fusobacterium* in our dataset and constituted 15.7% of the microbes found in the inflamed appendices (P: 6.8%, R: 2.0%), making it the genus with the highest relative abundance in the appendix samples.

**FIGURE 2.**
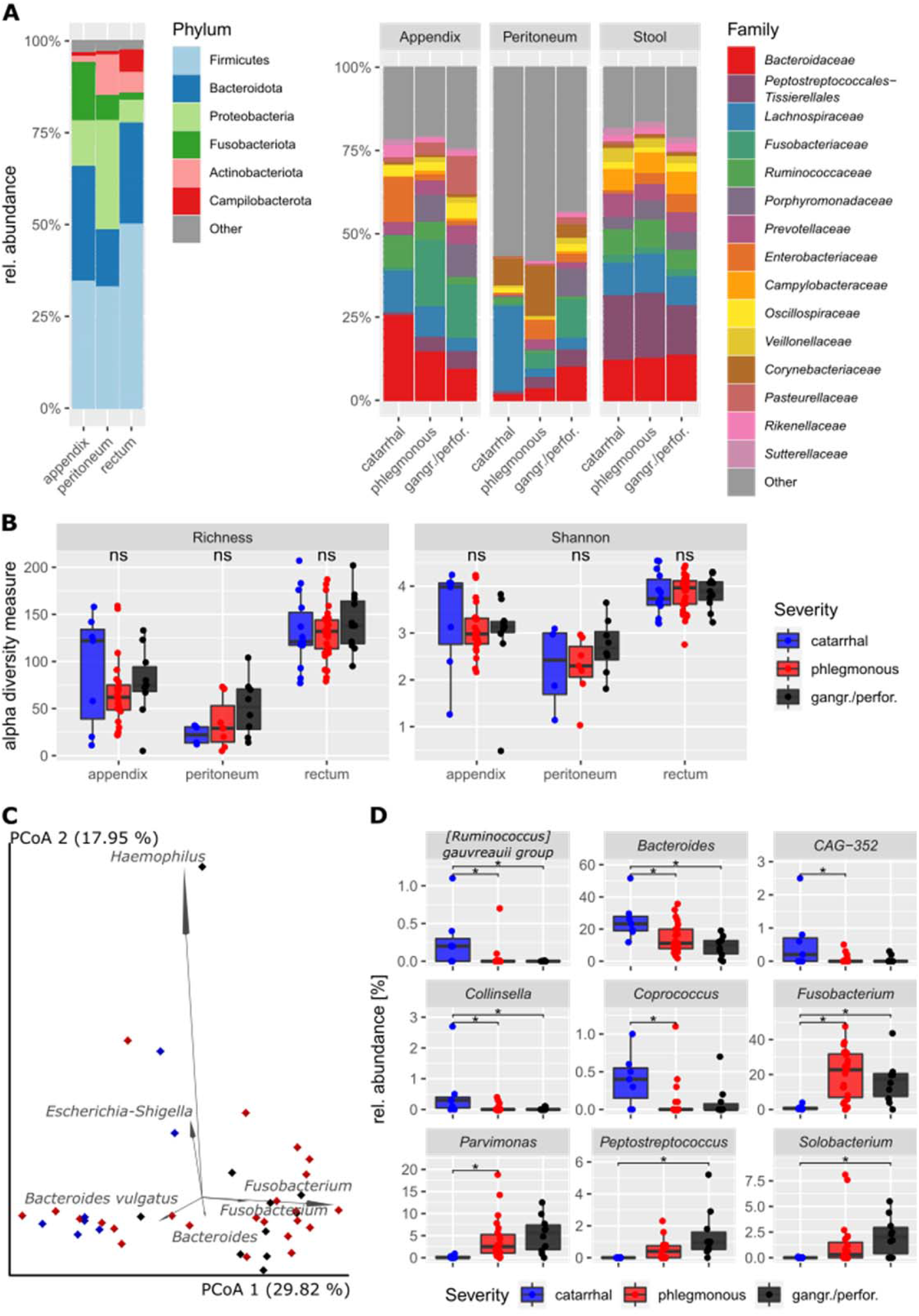
Bacterial diversity in children and adolescents with acute appendicitis (*n* = 60). **(A)** Microbial composition on phylum and family levels. **(B)** Alpha diversity measures (left, ASV richness; right, Shannon diversity) are not significantly different within the three sample types (pairwise Wilcoxon signed-rank test, all within-sample type comparisons *P*_adj_ > .26). For appendix samples **(C),** the biplot analysis of the six most important ASVs (weighted UniFrac PCoA) and **(D)** the nine genera that are significantly different between severity grades are shown, respectively (MaAsLin2; *, *P*_adj_ < .05). The catarrhal severity grade was used as a reference for the differential abundance analysis.

As expected, rectal samples showed the highest alpha diversity among the three sample types with an average richness of 132 ± 30 ASVs, followed by the appendix and peritoneum samples with 75 ± 41 and 37 ± 28 ASVs per sample, respectively. While the alpha diversity was significantly different between the three sample types (richness and Shannon diversity; ANOVA: *P* < .001), we did not observe significant changes in relation to disease severity (Fig. 2B). In terms of beta diversity, a significant difference from catarrhal to phlegmonous and from catarrhal to gangrenous/perforated appendicitis was observed in appendix samples (but not in rectal and peritoneal samples; weighted UniFrac and Bray-Curtis PERMANOVA: *P*_adj_ < .05 and *P*_adj_ < .01, respectively; Fig. 2C and Suppl. Fig. 1). This indicates that disease progression in AA is defined by a local microbial shift towards a dysbiotic and inflammation-promoting microbiota.

The results of the biplot analysis suggest that *Bacteroidetes* spp. and *Fusobacterium* spp. serve as indicators for disease severity in AA, as they tend to reflect catarrhal and gangrenous/perforated severity, respectively (Fig. 2C). Interestingly, two *Fusobacterium* ASVs were found among the six highest-impact features in the biplot analysis for both appendix and peritoneum samples. *Fusobacterium* has been repeatedly reported to be associated with disease severity in AA (12, 13, 15, 20, 23, 24), and the results of differential abundance analysis confirm this finding in our dataset (Fig. 2D). By applying MaAsLin2 (Multivariable Association Discovery in Population-scale Meta-omics Studies), we identified a total of nine differentially abundant genera, in which the abundances significantly increased or decreased upon disease progression in appendix samples. In addition to observing a significant increase in the relative abundance of *Fusobacterium* signatures, we also observed an increased abundance of other typically oral cavity-associated microbes including *Parvimonas*, *Peptostreptococcus* and *Solobacterium*. Strikingly, all these (potentially) opportunistic pathogens show a very low abundance at catarrhal severity (< 1% rel. abundance) and display a successive local expansion upon disease progression. In contrast, the respective abundance of the genera *Bacteroides*, *Ruminococcus gauvreauii* group, *Collinsella, Coprococcus* and *CAG-352* (family Ruminococcaceae) was found to be significantly increased in catarrhal appendicitis as compared with higher severity (Fig. 2D).

Surprisingly, we did not observe significant microbial differences in rectal and peritoneum samples that were associated with disease severity or any other clinical parameter. In rectal samples, we only observed certain tendencies, such as a successive increase in the genus *Escherichia-Shigella* from acute to gangrenous/perforated severity, based on the results of a pairwise Kruskal-Wallis test (*P* = .026); however, none of these results remained significant after FDR correction. Thus, we hypothesize that appendectomy has only limited impact on rectal samples. In contrast, peritoneum samples show a high heterogeneity between severity grades, displaying a high abundance of skin-associated microbes (e.g. *Staphylococcus:* 7.9%, *Streptococcus:* 10.3%, *Pseudomonas* 5.9%) in cases of catarrhal severity and shifting towards a microbial community that resembled the appendiceal microbiome in cases of phlegmonous and gangrenous/perforated severity. The apparent lack of significance of these results may be explained by the low number of catarrhal and phlegmonous samples with detectable bacterial signals (*n* = 4 and 7, respectively).

### Impact of archaea and fungi in acute appendicitis

In all sample types, archaeal taxa were dominated by the genera *Methanobrevibacter* (A: 90.6%, P: 78.4%, R: 69.5%) and *Methanosphaera* (A: 8.7%, P: 4.4%, R: 22.7%). Few samples yielded signatures of Methanomethylophilaceae or the thaumarchaeotal families Nitrososphaeraceae and *Candidatus Nitrosotenuis*, which were believed to represent skin contaminants. We did not observe significant differences between the archaeal composition and disease severity (or any other clinical parameter). Interestingly, *Methanosphaera* abundance and prevalence increased with AA severity, but not significantly (Kruskal-Wallis test for catarrhal and gangrenous/perforated severity: *P* = .08).

In ITS2 gene amplicon sequencing, only 12 appendix samples yielded detectable amounts of fungal signatures after quality control. *Malassezia restricta* represented the most prevalent fungal species that was predominantly found in the non-inflamed and catarrhal appendicitis samples. However, again no significant association between fungal taxa and disease severity or other clinical parameters was observed.

### Community state type analysis

As mentioned earlier in this communication, AA is hypothesized to comprise two different diseases: complicated appendicitis, which reaches the stage of perforation eventually, and uncomplicated appendicitis, which does not. In many cases, however, it is very difficult or even impossible to assess whether an unperforated appendix would have developed a perforation later on. As we observed a marked shift in the abundance of several bacterial taxa based on disease severity, we further investigated whether the microbial composition could be specific for complicated and uncomplicated appendicitis, respectively. In order to test this hypothesis, we performed a *de novo* community state type (CST) clustering based on the abundances of normalised ASV in the bacterial-dominated universal dataset.

Notably, we obtained a total of three different CSTs, two of which correlate well with the proposed disease types (Fig. 2). Within CST 2, most of the gangrenous/perforated and, in fact, all of the perforated cases cluster together. This CST is postulated to represent cases of complicated appendicitis, with the key taxa including oral-cavity-associated *Fusobacterium*, *Porphyromonas* and *Parvimonas* species (Fig. 2C) which, as we and others show (12, 13, 15, 23, 24), correlate closely with disease severity. CST 3, on the other hand, is believed to represent cases of uncomplicated appendicitis, as this group contains almost all cases of catarrhal appendicitis, with the gut-associated *Bacteroides* and *Faecalibacterium* as the dominant taxa. The major discriminating factor between these clusters is the disease severity, which is significantly higher in CST 2 as compared to CST 3 (chi-squared test, *P* = .004). This is also reflected in the number of patients with elevated blood leukocytes: 86.3% of CST 2 patients were diagnosed with leucocytosis but only 57.1% of CST 3 patients (*P* = .048). However, other diagnostic parameters did not show significant associations with both CSTs, including CRP (Wilcoxon rank-sum test; *P* = .470), PAS (*P* = .077) and Alvarado score (*P* = .083). Interestingly, we also observed another cluster of four samples in CST 1. This group is unique as we identified very high abundances of either *Haemophilus* (rel. abundance of 98.9% and 40.7%) or *Escherichia-Shigella* (rel. abundance of 28.9% and 53.5%) in the corresponding samples. CST 1 thus represents the “bacterial overgrowth” cluster, in which a single genus or species is hypothesized as being responsible for the AA.

### Metagenomic analysis of inflamed appendices

16S rRNA gene amplicon sequencing provides data about the microbial composition but cannot reliably be used to resolve taxa beyond the genus level or infer the genetic functionality of the microbiome. For this reason, we performed shotgun metagenomic sequencing of all 60 appendix samples and strove to identify key species/subspecies and virulence factors involved in AA pathogenesis. The results of the 16S rRNA gene analysis revealed the species *Fusobacterium necrophorum* (pairwise Kruskal-Wallis; *P* = .029), *Peptostreptococcus stromatis (P* = .042) and *Solobacterium moorei (P* = .042) to be enriched in gangrenous/perforated compared to catarrhal appendicitis. Furthermore, two *Porphyromonas* species (*P. uenonis, P* = .006 and *P. asaccharolytica, P* = .01) were also enriched in gangrenous/perforated samples. However, none of the enriched taxa were found to be significantly differently abundant after FDR correction. Such a lack of significant differences was also reported recently by Yuan et al. (25) who used a similar classification for AA severity.

Again, it is likely that pathological categorisation is not necessarily linked to the microbial composition alone, especially for phlegmonous cases. Therefore, we applied the 16S rRNA gene amplicon sequencing-based CST clustering method to our metagenomic data and performed a differential abundance analysis on both the taxonomic and functional levels. Based on MaAsLin2, only a significant enrichment of *Fusobacterium necrophorum* (*P*_adj_ = .035) was determined in CST 2 as compared to CST 3, further highlighting the importance of Fusobacteria with respect to disease severity. *Porphyromonas asaccharolytica* was also found to be enriched in CST 2 but not significantly after FDR correction (*P*_adj_ = .14). This microbial shift was accompanied by an altered abundance of functional genes in the community. In particular, we observed a higher abundance of catabolism pathways in CST 2 and especially for amino acids, including lysine fermentation to crotonyl-CoA (*P*_adj_ = .035), histidine degradation (*P*_adj_ = .051), glutamate fermentation (*P*_adj_ = .066) and the associated Na-driven 2-hydroxyglutarate pathway (*P*_adj_ = .051), as well as the bacterial proteasome pathway (*P*_adj_ = .051).

To further validate and test the robustness of the results reported above, DESeq2 was performed (Fig. 3). On the taxonomic level, *F. necrophorum* was confirmed as being significantly enriched in more complicated cases (CST 2) but also *F. nucleatum*, two *Porphyromonas* species (*P. endodontalis* and *P. uenonis*) and two unspecified species of the genera *Prevotella* and *Alloprevotella* were significantly enriched in CST 2. Despite the marked expansion of those species, no significant change was observed at the functional level. As noted in the MaAsLin2 analysis, an enrichment of catabolic pathways was apparent in CST 2, indicating a potentially increased release of nutrients by, e.g. apoptotic or necrotic host cells. However, this hypothesis is highly speculative and needs to be verified via physiological characterisation of the corresponding species.

**FIGURE 3.**
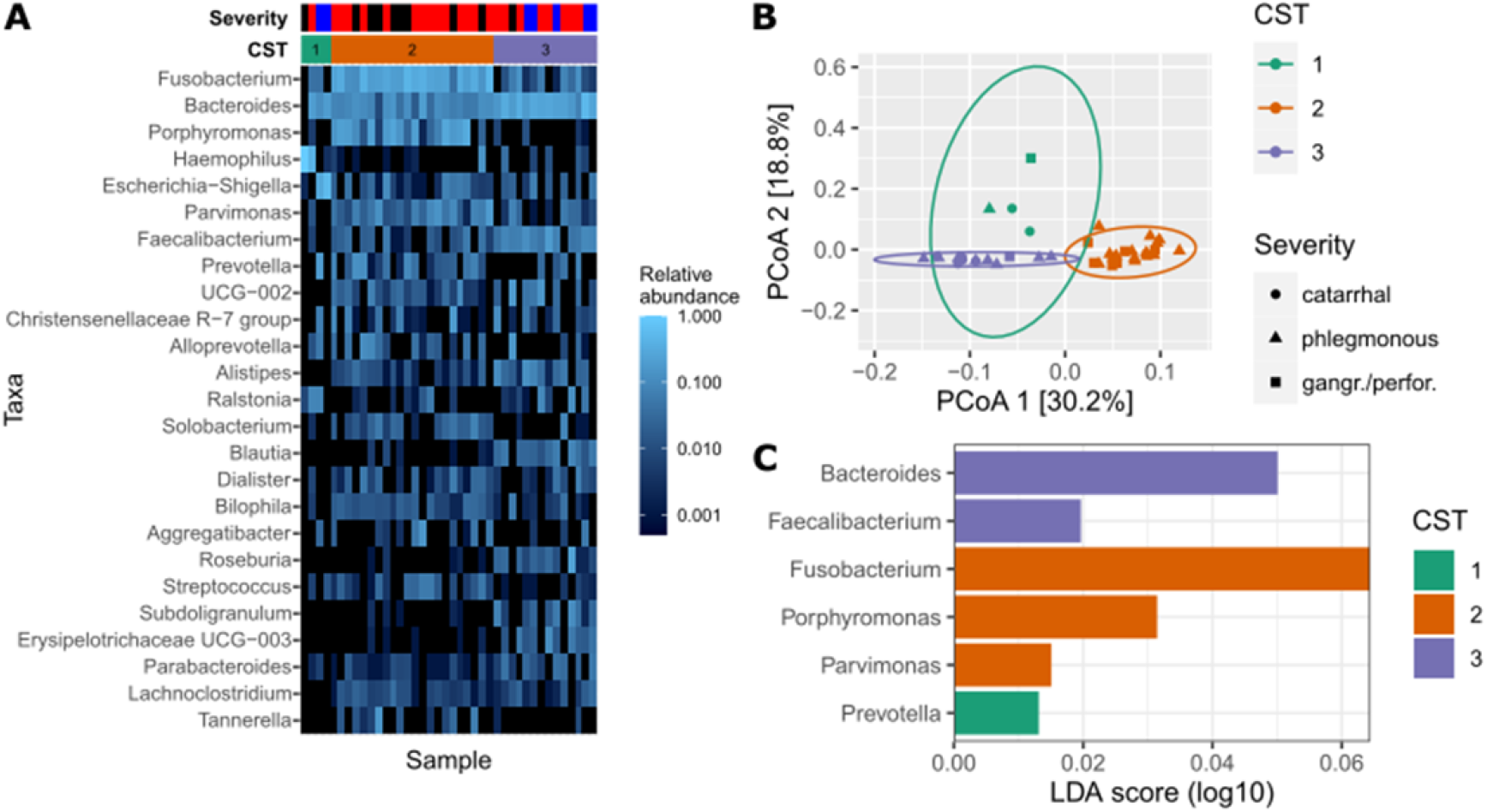
Community state type (CST) analysis for appendix samples. **(A)** shows a heatmap of the 25 most abundant genera in the appendix samples, sorted by the three defined CSTs. Appendicitis severity grades are as follows: blue, catarrhal; red, phlegmonous; black, gangrenous/perforated. **(B)** CST clustering is shown and based on Principal Coordinates Analysis (PCoA). **(C)** Indicates the most important genera that contribute to the corresponding CSTs as defined by Linear discriminant analysis Effect Size (LEfSe) analysis.

Neither the differential abundance nor ABRicate analysis yielded any indications that antimicrobial resistance or virulence genes were enriched among the disease severities.

## Discussion

In this study, we identify microbes that are characteristic and potentially responsible for the different severity stages of AA. We observed a shift in the microbial community in the appendix (Fig. 2), depending on the severity of the disease, that was primarily attributed to a change in the bacterial beta diversity, but not to an increased richness, as previously suggested (20). This observed shift in diversity is almost exclusively due to changes in bacterial taxa without any significant contribution from fungi or archaea. Both 16S rRNA gene amplicon and metagenomic shotgun sequencing of the appendix microbiome clearly indicate a local expansion of mainly oral cavity-associated microbes in complicated appendicitis, including *Fusobacterium*, *Porphyromonas*, *Parvimonas*, *Peptostreptococcus* and *Solobacterium*. Likewise, we could provide support for the previous observation of a stepwise decline in gut-associated *Bacteroides* as the disease severity increased (13, 20, 23, 24). This decline is accompanied by a substantial reduction in the relative abundance of *Ruminococcaceae*, *Collinsella* and *Coprococcus* from catarrhal to phlegmonous and/or gangrenous/perforated appendicitis (Fig. 2D).

For over a decade now, researchers have reported that uncomplicated and complicated appendicitis do not share the same etiopathogenesis. This claim is supported by the observation – among others – that not all cases of AA lead eventually to perforation. While the time span from pain onset to surgery positively correlates with perforation, some cases remain phlegmonous even after a long duration of pain (61–63). Our data on the microbial composition support this difference in etiopathogenesis for AA, as we observed a decisive microbial shift from catarrhal to gangrenous/perforated appendicitis, indicated by the results of the differential abundance and beta diversity analyses as well as by *de novo* CST analysis (Fig. 3). We identified three CST clusters with different characteristics. The first cluster is associated with bacterial overgrowth typified by either *Haemophilus* or *Escherichia-Shigella* (CST 1). The second cluster was enriched with Fusobacteria and other oral cavity-associated microbes (such as *Porphyromonas* and *Parvimonas*), a characteristic which appears to be a hallmark of complicated appendicitis (CST 2). In the third cluster, cases are defined by higher relative abundance of typical gut-associated bacteria such as *Bacteroides* and *Faecalibacterium* without an apparent enrichment of oral microbes. These cases might be unlikely to develop perforation and can be attributed to uncomplicated AA. However, while CST 3 and CST 2 clearly separate catarrhal and perforated appendicitis, the phlegmonous and (to a minor extent) gangrenous cases are distributed in both clusters. It is plausible that the microbial community is only partially responsible for AA severity and that some patients may develop complicated appendectomy with a CST-3-like microbial composition. AA is a multifactorial disease that depends on multiple aspects including lifestyle, diet and genetic predisposition (1, 2, 4, 16). It is apparent that proper tissue function and microbial homeostasis require a delicate balance between the immune system, microbiome and host epithelial cells. Previous work by Rivera-Chavez *et al*. (16) for example, indicated that single nucleotide polymorphisms in the IL-6 gene can partially explain the development of complicated appendicitis. Thus, an analysis of the microbiome alone does not allow us to explain the disease etiopathogenesis and an uncomplicated microbial state (CST 3) may still (rarely) lead to gangrene. Due to a lack of longitudinal data, we also cannot exclude the possibility that CSTs can shift from one state to another as the disease progresses. For example, uncomplicated (CST 3) may shift to complicated appendicitis (CST 2) upon the further growth of Fusobacteria and colonisation by *Porphyromonas* and *Parvimonas* species.

The latest and current findings indicate that special attention should be paid to the presence of Fusobacteria in appendicitis patients, especially when other oral pathogens such as *Porphyromonas* or *Parvimonas* are present. Fusobacteria are robustly and frequently associated with an increased disease severity and appear to be hallmark taxa in the development of complicated appendicitis. While species of this taxon were also found to be part of the normal appendix microbiota (12, 24), their abundance, prevalence and tissue invasion are greatly increased in complicated appendicitis (Figs. 2 and 3) (23). The pathogenic expansion and tissue invasion in AA is mainly attributed to the species *F. nucleatum* and *F. necrophorum* (Fig. 4) (23, 64), two well-known, opportunistic pathogens that are also associated with other gastrointestinal diseases such as inflammatory bowel disease, primary sclerosing cholangitis and colorectal cancer (65). *Fusobacterium nucleatum* is a mutualistic microorganism that interacts with human tissue in ways that range from neutral to pathogenic. Several disease-promoting mechanisms have been described, ranging from immunomodulatory effects to tissue and cell invasion and on to recruitment and virulence enhancement of other, potentially pathogenic microbes (nicely reviewed in Brennan and Garrett (66)). As such, *F. nucleatum* has been shown to induce antimicrobial and pro-inflammatory host responses (such as β-defensin, IL-6 and IL-8 expression) and to be capable of actively penetrating and surviving in human tissue and immune cells. Furthermore, Fusobacteria are important biofilm-forming bacteria. Due to the high variety of adhesins they produce, *F. nucleatum* can bind to host cells, enabling it to act as a docking hub for other microorganisms due to its elongated shape. In colorectal cancer, Fusobacteria frequently co-occur with other oral microbes such as *Peptostreptococcus* spp. and *Leptotrichia* spp. (67, 68) and it is feasible that a *Fusobacterium*-mediated colonisation of secondary oral pathogens such as *Porphyromonas*, *Parvimonas*, *Solobacterium* or *Peptostreptococcus* also occurs in complicated appendicitis. All of the mechanisms employed by Fusobacteria are common virulence factors that aid bacteria in establishing new niches, acquiring nutrients and evasion of the immune system (69). As such, the increase in the relative abundance of amino acid metabolising pathways that was observed in the metagenomic data may be directly linked to the growth of Fusobacteria. Free lysine, histidine, glutamate and serine are required for *F. nucleatum* growth (70), and the relative abundance of the catabolic pathways for the production of the former three amino acids were enhanced in CST 2. Especially lysine fermentation to crotonyl-CoA was predominantly found to be enriched in more severe cases (CST 2), and this pathway has been described in only few microbes, including *F. nucleatum* and *Porphyromonas gingivalis* (71). It is tempting to speculate that Fusobacteria actively trigger apoptosis in intestinal epithelial cells during AA to release peptides and amino acids.

**FIGURE 4.**
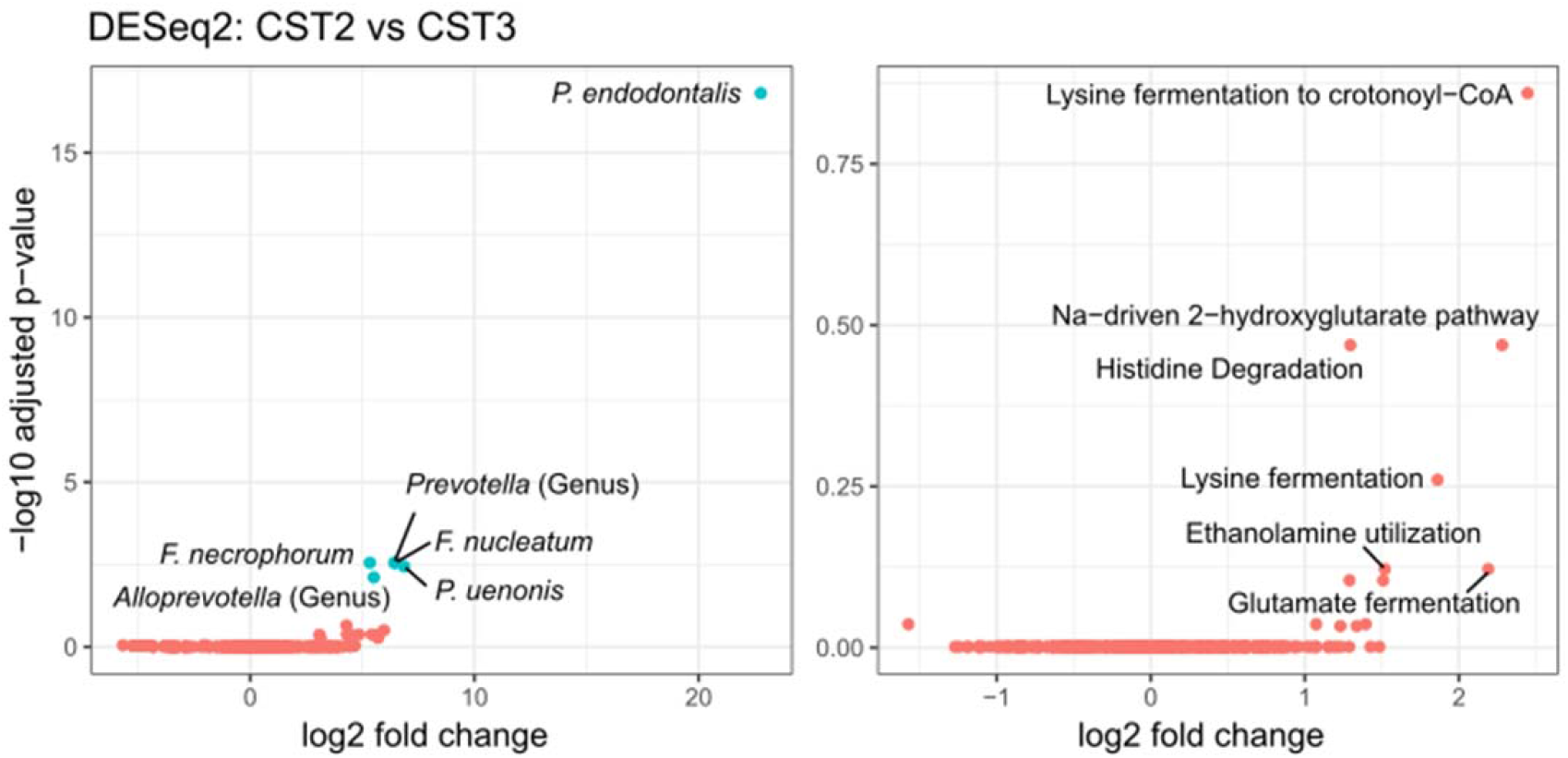
DESeq2 analysis comparing community state types (CST) 2 and 3 on taxonomic (left) and functional level (right). In both plots, the six most important features were labelled and the significantly differentially abundant features are highlighted in blue (*P_adj_* < .05 and fold change > 1.5). Species abbreviations: *Fusobacterium necrophorum, Fusobacterium nucleatum, Porphyromonas endodontalis*, *Porphyromonas uenonis*.

Considering all of these aspects, Fusobacteria appear to play a causative role in several diseases. However, other researchers have also suggested that Fusobacteria only plays a passenger role in disease (Tjalsma 2012), especially since not enough evidence is available to satisfy Koch’s postulates. The same holds true in case of AA. Fusobacteria are certainly associated with disease severity, and particularly with complicated appendicitis, but we do not yet know whether they play a role as a driver or passenger in the disease.

In recent decades, major improvements in AA diagnosis have been achieved, e.g. by implementing and evaluating standardised clinical scoring systems, such as PAS and Alvarado, or by applying routine imaging techniques to the lower abdomen, including ultrasound or computed tomography (72). These tools help physicians choose the appropriate therapy and can help to prevent unnecessary surgery. Reliable and easily accessible diagnostic and microbial markers could help to further improve AA diagnosis and even might indicate whether antibiotic treatment is suitable.

Unfortunately, we did not detect significant differences in the microbiome of rectal samples within our study cohort. The local expansion of opportunistic pathogens within the appendix had no apparent effect on the rectal microbiome. This result may arguably be due to the high diversity of microbes in the large intestine along with a “dilution” of signatures or microbes from the appendix as they pass through the large intestine. Rectal samples are not suitable for distinguishing complicated and uncomplicated appendicitis but may be useful for discriminating between AA and healthy patients or patients with other gastrointestinal diseases that cause abdominal pain, such as colitis. In fact, previous studies have suggested that rectal samples from AA patients show an elevated richness and increase in the abundance of *Bulleidia*, *Dialister* and *Porphyromonas* as compared to healthy controls (20). Longitudinal analyses prior to as well as after appendectomy are required to show whether microbial changes in the distal colon are causally linked with AA.

In peritoneal samples, we observed a stepwise but not significant shift from skin-associated taxa dominating in catarrhal cases to intestinal-associated taxa dominating in complicated appendicitis cases. In fact, peritoneal samples may serve as a proxy for AA severity, as invasive pathogens such as *Fusobacteria* can be detected both in patients with perforated appendicitis and even in some cases with phlegmonous and gangrenous appendicitis. While the diagnostic value of peritoneal samples is rather low since sampling requires invasive interventions, we could confirm the presence of pathogen signatures in the peritoneal area even in non-perforated appendicitis. The health implications of this finding are uncertain but underline the importance of quick therapeutic intervention in case of AA via surgery or antibiotics.

Furthermore, we observed a significantly higher incidence of leucocytosis in complicated as compared to uncomplicated appendicitis cases. While the diagnostic value of white blood cell and leukocyte counts in AA is controversial (73), the presence of “unknown” microbes in the GIT, such as oral cavity-associated pathogens, may be indicated by increased leukocytes counts.

Several questions regarding the etiopathogenesis of AA are still open. First of all, the origins of appendicitis are still unknown. One possibility was proposed by A. Swidsinsky and his team (23), who postulated that appendicitis is a pathogen-induced infectious disease. This idea was discussed recently in the context of the COVID-19 crisis, as the number of paediatric AA cases substantially dropped (~40%) in 2020 as compared to those in previous years (74, 75). Even though the reasons for this development are unknown, it is possible that social distancing and improved hygiene could have contributed to lower incidence rates by reducing exposure to specific microbes. Furthermore, a recent analysis of stimulated peripheral blood mononuclear cells did not show evidence for innate immune dysfunction in patients with a history in AA, but suggested a possible connection regarding responses between complicated and uncomplicated appendicitis (18). These findings further strengthen the idea that bacterial infection and/or dysbiosis causes AA. In this regard, it is also of crucial importance to trace the origin of the potential pathogens. Recently, the impact of oral microbes in several inflammatory gastrointestinal diseases was highlighted and received an increased amount of attention (21, 65, 76, 77). The history of dental health and oral cavity integrity in AA patients should be investigated in further studies as well, since both factors may show a putative link to appendicitis severity.

### Limitations

AA is a multifactorial disease defined by several host- and microbiome-based factors. Subsequent studies should not be restricted to a singular aspect of this disease, but include holistic analyses of both the host (genetics, immunological parameters) and microbiome (composition, metabolomics, transcriptomics). Considering the potentially negative effects of oral pathogens involved in AA, saliva and subgingival sampling should also be considered in future studies. Another limitation was the inclusion of only 60 patients in our study. Whether the tendencies and borderline-significant results we show are truly associated with AA could be further resolved in studies with larger cohorts, ideally including a suitable number of incidental appendectomies that could serve as controls.

Furthermore, one-third of the patients were treated with antibiotics during or immediately before surgery. As discussed earlier, we did not expect an effect on the microbial composition, but we also cannot fully exclude it. Last, our metagenomic sequencing was limited in terms of the overabundance of host-mapped sequences, restricting the analysis to the gene-centric approach described (see Material and Methods). Further studies need to be carried out to optimise sampling and/or sample processing to reduce the amount of human DNA for microbial analysis.

### Conclusion

We show that uncomplicated AA was characterised rather by the increased relative abundance of typical gastrointestinal microorganisms, whereas complicated AA was associated with misplaced oral microorganisms. Fusobacterium and its associated fatal group are obviously involved in necrotizing activities, which could substantially lead to barrier breaks and perforation. As these microorganisms originate from the oral cavity, their transmission into the gastrointestinal tract needs to be resolved further, including an examination of the function of the stomach barrier or dental issues. Although a reliable microbiome-based biomarker for distinguishing uncomplicated and complicated AA could not be identified in easily-accessible samples in this study, other (clinical) parameters could fulfil this requirement. As a consequence, further studies are required to delineate the clinical phenotype of *Fusobacterium* infection in appendicitis.

## Materials and Methods

### Study design

A total of 60 children and adolescents with acute appendicitis (AA) undergoing appendectomy at the Department of Paediatric and Adolescent Surgery at the Medical University of Graz from April to June 2019 were prospectively recruited. The study was approved by the Ethics committee of the Medical University in Graz (31-004 ex 18/19) and performed according to the Helsinki Declaration. Written informed consent was given by the participants and/or the caregivers after they were provided with written and oral information about the study. All data and samples were pseudo-anonymised.

### Sample and data collection

Preoperatively, demographic data and serum parameters were obtained, including total leukocyte count and C-reactive protein (CRP). Additionally, the Alvarado score (27) and Pediatric Appendicitis Score (PAS) (28) were assessed.

Rectal (*n* = 60) and in cases of fluid collection in the abdominal cavity peritoneal swabs (*n* = 34) were taken with sterile nylon swabs (FLOQSwabsTM, Copan) prior to and during the operation, respectively. After the removal of the vermiform appendix, about 1 cm of the proximal part was excised and stored at −80°C together with the other samples until further processing. The remaining tissue was used for standard histological examination. According to Carr (26), catarrhal appendicitis is defined as a local inflammation with few intraepithelial neutrophils and reactive intraepithelial changes. Phlegmonous appendicitis was diagnosed in cases with evidence of neutrophils invading the mucosa, submucosa and muscularis propria, intraluminal abscess and invasion of the surrounding tissue. Gangrenous appendicitis additionally showed intramural necrosis. Perforation was defined as microscopic or macroscopic perforation in the abdominal cavity.

### Sample processing

Appendix samples were processed as follows: While keeping the tissue deep-frozen, two cross sections of about 2-3 mm were cut with a sterile scalpel. The first slice was discarded, and about 25 mg tissue were collected from the second, making sure to include luminal content. DNA was extracted from tissue and rectal/peritoneal swabs using the DNeasy^®^ PowerSoil^®^ Kit (Qiagen-Hilden) according to manufacturer’s instructions with the following exception: Lysis and homogenisation of the samples was performed in two cycles of bead beating using a MagNA Lyser (Roche Diagnostics GmbH) at 6500 rpm for 30s with intermediate cooling. The DNA concentration was measured using the Qubit™ dsDNA HS Assay Kit (ThermoFisher Scientific).

### PCR and amplicon sequencing

To identify the microbial communities from the appendix, rectum and peritoneum, we amplified the V4 region of the 16S rRNA gene to detect bacteria and archaea and the internal transcribed spacer 2 (ITS2) region of the 23S rRNA gene to detect fungi. Sequences from prokaryotic specimens were amplified with the universal primer pair 515F and 806R containing Illumina adapter sequences (29). We performed a nested PCR to specifically amplify archaea DNA by using the primer combination 344F-1041R/Illu519F-Illu806R as described previously (30). PCR reactions were carried out in a final volume of 25 μL containing: TAKARA Ex Taq® buffer with MgCl2 (10 X; Takara Bio Inc.), primers (200 nM of each), dNTP mix (200 μM of each), TAKARA Ex Taq® Polymerase 0.5 U, water (Lichrosolv®; Merck) and DNA template (1-2 μL of genomic DNA). To amplify DNA from fungal specimens, we used the primer pair ITS86F-ITS4 containing Illumina adapter sequences (31). The PCR reaction was carried out with the same setup as described above, but 400 nM of each primer were used instead. All primers and PCR conditions used are listed in Tables S1 and S2.

Both the library preparation and sequencing of the amplicons were performed at the Core Facility Molecular Biology, Center for Medical Research at the Medical University Graz, Austria. In brief, SequalPrep™ normalisation plates (Invitrogen) were used to normalise the DNA concentration, and each sample was subsequently indexed with a unique barcode sequence (8 cycles index PCR). All indexed samples were pooled, and the products of the indexing PCR were purified with gel electrophoresis. Sequencing was performed using an Illumina MiSeq device and the MS-102-3003 MiSeq® Reagent Kit v3-600 cycles (2×251 cycles).

The MiSeq data for all three approaches (universal, archaeal, fungal) were analysed individually using QIIME2 V2019.11 (32) as described previously (33). Briefly, the DADA2 algorithm (34) was used to demultiplex and de-noise truncated reads as well as to generate amplicon sequence variants (ASVs). Taxonomic assignment was based on the SILVA v138 database (35) for prokaryotic specimens and on the UNITE v8.3 database (36) for fungi. Fungal raw reads were also pre-processed with ITSxpress (37), trimming reads to the desired ITS2 region. The datasets was filtered as follows: Potential contaminants were identified and removed with the R software package decontam (38) by providing negative controls (DNA extraction and PCR negative controls) and applying a threshold of 0.25. Control samples were subsequently removed from the dataset. Unassigned ASVs, those classified as chloroplast and mitochondria, and ASVs with fewer than 10 total reads were also removed. Rarefaction of the datasets was performed by scaling with ranked subsampling (SRS) (39) using rarefaction depths of 1000, 100 and 50 for bacteria, fungi and archaea, respectively.

### Whole Genome Sequencing (WGS)

Shotgun metagenome sequencing was performed for all 60 appendix samples. A total of 200 ng of extracted DNA from each sample was sent to Macrogen Europe (Amsterdam, Netherlands). Library extraction was performed with the TruSeq DNA PCR-Free kit (Illumina) and sequenced with the Illumina NovaSeq 6000 platform (Illumina).

Raw reads were processed as described previously (33). In brief, quality control and filtering were performed with fastqc (v0.11.8) (40) and trimmomatic (v0.38) (41). Accepted reads were mapped to the human chromosome hg19 with bowtie2 (v2.3.5) (42), and unmapped reads were retained using samtools (v1.9) (43). Bedtools (v2.29.0) (44) was subsequently used to extract host-removed files. Unfortunately, a genome-centric approach was not possible for this dataset, as the assembly with Megahit (v1.1.3) (45) and the subsequent binning using MaxBin (v2.2.4) (46) did not yield any draft genomes. Even though all samples had ~20 Mil. raw reads, about 99% of all reads mapped to the human genome. This high abundance of host reads could not be mitigated by using the NEBNext® Microbiome DNA Enrichment Kit in our approach. Instead, we performed a gene-centric analysis, performing blastx with diamond (v2.0.8) (47) to annotate the host-removed, quality-filtered reads against the NCBI nr database (release Sep. 2020). Megan (6.18.0) (48) was used to remove reads that were classified as Metazoa and to compare samples based on normalised counts. Subsequently, the taxonomic and functional annotation based on the SEED database (49) was exported and used for statistical analysis.

In addition, contigs were screened for antimicrobial resistance and virulence genes with ABRicate using all available databases: ARG-ANNOT, CARD, EcOH, Ecoli_VF, MEGARES 2.00, NCBI AMRFinderPlus, PlasmidFinder, Resfinder and VFDB (50). Furthermore, we mapped the quality filtered contigs against the genome of *Fusobacterium nucleatum* (NCBI Reference Sequence: NZ_LN831027.1) and several archaeal genomes, using species that were repeatedly reported to be associated to the human gut, including *Methanobrevibacter smithii* (DSM 2374 and strain WWM1085), *Methanosphaera stadtmanae* (DSM 3091), *Methanomassiliicoccus luminyensis* (B10), *Methanocorpusculum*, *Methanobacterium*, *Halorubrum lipolyticum* (DSM 21995) and *Haloferax* sp. Arc-Hr (51).

### Statistical analysis

Statistical analysis of clinical parameters was performed using the IBM® SPSS® Statistics 26 software package. After testing for Gaussian normal distribution using the Shapiro-Wilk test, Kruskal-Wallis tests were applied to nonparametric data, while the comparison of parametric data was conducted with a one-way analysis of variance (ANOVA) and a Tukey post-hoc test. Comparisons of categorical data were performed with the chi-square test. Microbial data analysis and visualization were done with QIIME2 (V2019.11) and R (V4.1.1). Phylogenetic distances for the amplicon sequencing data were calculated in QIIME2 using the fasttree plugin (52) and subsequently analysed following the core-metrics-phylogenetic command without further subsampling. Biplots were performed based on the weighted UniFrac distance matrix and were calculated using the biplot plugin in QIIME2. The Emperor plugin (53, 54) was used to illustrate the results of the biplot analysis.

Differential abundance and alpha diversity testing as well as visualisation were performed using normalised data within R and the packages phyloseq (55), ggplot2 (56), Maaslin2 (57), DESeq2 (58). A full list of all packages and versions used can be found in Table S3. Alpha diversity was calculated based on the filtered and normalised dataset, and mean values ± standard deviation (SD) are reported. After testing for normality, pairwise Wilcoxon signed-rank tests were performed, and the resulting *P*-values were corrected for false discovery rates (FDR) according to Benjamini and Hochberg (59). For MaAsLin2, the default settings were applied except for max_significance = 0.05 and min_prevalence = 0.001. DESeq2 analysis was considered significant if *P*_adj_ < 0.05 and log2FoldChange ≥ 0.58 (corresponds to a 1.5-fold change).

### Community state type (CST) analysis

We performed *de novo* community state type (CST) clustering based on rarefied 16S rRNA gene data (universal primers) and the corresponding weighted UniFrac distance matrix. Calculation of the CSTs was applied using the methodology of DiGiulio et al. (60). Based on gap statistics, the number of clusters was determined (*k* = 3).

## Data availability

The datasets supporting the conclusions of this article are available from the European Nucleotide Archive (ENA) repository, Primary Accession: PRJEB49215 in https://www.ebi.ac.uk/.

## Acknowledgements

The work was funded by the Austrian Science Fund FWF, no. P32697, whose support is gratefully acknowledged. M.B. is a student in the local PhD doctoral programme “MolMed”. The authors acknowledge the support of the ZMF Galaxy Team: Core Facility Computational Bioanalytics, Medical University of Graz, funded by the Austrian Federal Ministry of Education, Science and Research, Hochschulraum-Strukturmittel 2016 grant as part of BioTechMed Graz.

We also acknowledge the support and cooperation of all participants, doctors and nurses in the Department of Paediatric and Adolescent Surgery in Graz who contributed to this study. Special thanks to Valentin Trinkl for his support in sample preparation. We appreciate the help of Christina Kumpitsch, Lisa Wink, Viktoria Weinberger and Charlotte Neumann for critical proofreading and Gregor Gorkiewicz for his scientific advice.

Authors contributed as follows. G.S., C.M.-E.: study design. G.S., C.F., C.C., H.T.: sampling, collection of metadata. M.B., K.B.: biological samples processing. M.B.: microbiome data and statistical analysis, preparation of figures and tables, wrote manuscript. C.M.-E., A.M.: support of microbiome data and statistical analysis. All authors: supported manuscript writing and scientific contributions.

The authors declare that they have no competing interests.

## Supplementary files legends

**FIGURE S1 |** Weighted UniFrac PCoA Biplots of the six most important features for the corresponding sample sites.

**TABLE S1 |** Primer pairs used for archaeal, bacterial and fungal PCR.

**TABLE S2 |** PCR conditions for the three investigated approaches.

**TABLE S3 |** R plugins and versions used.

## References

1. Addiss DG, Shaffer N, Fowler BS, Tauxe R V. 1990. The epidemiology of appendicitis and appendectomy in the united states. Am J Epidemiol 132:910–925.

2. Lee JH, Park YS, Choi JS. 2010. The Epidemiology of Appendicitis and Appendectomy in South Korea: National Registry Data. J Epidemiol 20:97–105.

3. Ahmed HO, Muhedin R, Boujan A, Aziz AHS, Abdulla A muhamad, Hardi RA, Abdulla AA, Sidiq TA. 2020. A five-year longitudinal observational study in morbidity and mortality of negative appendectomy in Sulaimani teaching Hospital/Kurdistan Region/Iraq. Sci Rep 10:1–7.

4. Golz RA, Flum DR, Sanchez SE, Liu XH, Donovan C, Drake FT. 2020. Geographic Association between Incidence of Acute Appendicitis and Socioeconomic Status. JAMA Surg 155:330–338.

5. Wangensteen OH, Dennis C. 1939. EXPERIMENTAL PROOF OF THE OBSTRUCTIVE ORIGIN OF APPENDICITIS IN MAN. Ann Surg 110:629–47.

6. Singh J, Mariadason J. 2013. Role of the faecolith in modern-day appendicitis. Ann R Coll Surg Engl 95:48–51.

7. Arnbjornsson E, Bengmark S. 1983. Obstruction of the appendix lumen in relation to pathogenesis of acute appendicitis. Acta Chir Scand 149:789–791.

8. Luckmann R. 1989. Incidence and case fatality rates for acute appendicitis in California: A population-based study of the effects of age. Am J Epidemiol 129:905–918.

9. Luckmann R, Davis P. 1991. The epidemiology of acute appendicitis in california: Racial, gender, and seasonal variation. Epidemiology 2:323–330.

10. Salminen P. 2020. Acute Appendicitis Incidence-Predisposing Factors, From Microbiota to Socioeconomic Status? JAMA Surg 155:338–339.

11. Mariage M, Sabbagh C, Grelpois G, Prevot F, Darmon I, Regimbeau J-M. 2019. Surgeon’s Definition of Complicated Appendicitis: A Prospective Video Survey Study. Euroasian J hepato-gastroenterology 9:1–4.

12. Guinane CM, Tadrous A, Fouhy F, Anthony Ryan C, Dempsey EM, Murphy B, Andrews E, Cotter PD, Stanton C, Paul Ross R. 2013. Microbial composition of human appendices from patients following appendectomy. MBio 4:1–6.

13. Zhong D, Brower-Sinning R, Firek B, Morowitz MJ. 2014. Acute appendicitis in children is associated with an abundance of bacteria from the phylum Fusobacteria. J Pediatr Surg 49:441–446.

14. Peeters T, Penders J, Smeekens SP, Galazzo G, Houben B, Netea MG, Savelkoul PH, Gyssens IC. 2019. The fecal and mucosal microbiome in acute appendicitis patients: an observational study. Future Microbiol 14:111–127.

15. Rogers MB, Brower-Sinning R, Firek B, Zhong D, Morowitz MJ. 2016. Acute Appendicitis in Children Is Associated With a Local Expansion of Fusobacteria. Clin Infect Dis 63:71–78.

16. Rivera-Chavez FA, Peters-Hybki DL, Barber RC, Lindberg GM, Jialal I, Munford RS, O’Keefe GE. 2004. Innate Immunity Genes Influence the Severity of Acute Appendicitis. Ann Surg 240:269.

17. Rubér M, Berg A, Ekerfelt C, Olaison G, Andersson RE. 2006. Different cytokine profiles in patients with a history of gangrenous or phlegmonous appendicitis. Clin Exp Immunol 143:117–24.

18. Peeters T, Martens S, D’Onofrio V, Stappers MHT, van der Hilst JCH, Houben B, Achten R, Joosten LAB, Gyssens IC. 2020. An observational study of innate immune responses in patients with acute appendicitis. Sci Rep 10:17352.

19. Podda M, Gerardi C, Cillara N, Fearnhead N, Gomes CA, Birindelli A, Mulliri A, Davies RJ, Di Saverio S. 2019. Antibiotic treatment and appendectomy for uncomplicated acute appendicitis in adults and children: A systematic review and meta-analysis. Ann Surg 270:1028–1040.

20. Jackson HT, Mongodin EF, Davenport KP, Fraser CM, Sandler AD, Zeichner SL. 2014. Culture-Independent Evaluation of the Appendix and Rectum Microbiomes in Children with and without Appendicitis. PLoS One 9:e95414.

21. Blod C, Schlichting N, Schülin S, Suttkus A, Peukert N, Stingu CS, Hirsch C, Elger W, Lacher M, Bühligen U, Mayer S. 2018. The oral microbiome—the relevant reservoir for acute pediatric appendicitis? Int J Colorectal Dis 33:209–218.

22. Oh SJ, Pimentel MM, Leite GGS, Celly S, Villanueva-Millan MJ, Lacsina I, Chuang B, Parodi G, Morales W, Weitsman S, Singer-Englar T, Barlow GM, Zhai J, Pichestshote N, Rezaie A, Mathur R, Pimentel MM. 2020. Acute appendicitis is associated with appendiceal microbiome changes including elevated Campylobacter jejuni levels. BMJ Open Gastroenterol 7:1–10.

23. Swidsinski A, Dörffel Y, Loening-Baucke V, Theissig F, Rückert JC, Ismail M, Rau WA, Gaschler D, Weizenegger M, Kühn S, Schilling J, Dörffel W V. 2011. Acute appendicitis is characterised by local invasion with Fusobacterium nucleatum/necrophorum. Gut 60:34–40.

24. Salö M, Marungruang N, Roth B, Sundberg T, Stenström P, Arnbjörnsson E, Fåk F, Ohlsson B. 2017. Evaluation of the microbiome in children’s appendicitis. Int J Colorectal Dis 32:19–28.

25. Yuan J, Li W, Qiu E, Han S, Li Z. 2021. Metagenomic NGS optimizes the use of antibiotics in appendicitis patients: Bacterial culture is not suitable as the only guidance. Am J Transl Res 13:3010–3021.

26. Carr NJ. 2000. The pathology of acute appendicitis. Ann Diagn Pathol 4:46–58.

27. Alvarado A. 1986. A practical score for the early diagnosis of acute appendicitis. Ann Emerg Med 15:557–64.

28. Samuel M. 2002. Pediatric appendicitis score. J Pediatr Surg 37:877–81.

29. Walters W, Hyde ER, Berg-Lyons D, Ackermann G, Humphrey G, Parada A, Gilbert JA, Jansson JK, Caporaso JG, Fuhrman JA, Apprill A, Knight R. 2016. Improved Bacterial 16S rRNA Gene (V4 and V4-5) and Fungal Internal Transcribed Spacer Marker Gene Primers for Microbial Community Surveys. mSystems 1.

30. Pausan MR, Csorba C, Singer G, Till H, Schöpf V, Santigli E, Klug B, Högenauer C, Blohs M, Moissl-Eichinger C. 2019. Exploring the Archaeome: Detection of Archaeal Signatures in the Human Body. Front Microbiol 10:2796.

31. Op De Beeck M, Lievens B, Busschaert P, Declerck S, Vangronsveld J, Colpaert J V. 2014. Comparison and validation of some ITS primer pairs useful for fungal metabarcoding studies. PLoS One 9:e97629.

32. Bolyen E, Rideout JR, Dillon MR, Bokulich NA, Abnet CC, Al-Ghalith GA, Alexander H, Alm EJ, Arumugam M, Asnicar F, Bai Y, Bisanz JE, Bittinger K, Brejnrod A, Brislawn CJ, Brown CT, Callahan BJ, Caraballo-Rodríguez AM, Chase J, Cope EK, Da Silva R, Diener C, Dorrestein PC, Douglas GM, Durall DM, Duvallet C, Edwardson CF, Ernst M, Estaki M, Fouquier J, Gauglitz JM, Gibbons SM, Gibson DL, Gonzalez A, Gorlick K, Guo J, Hillmann B, Holmes S, Holste H, Huttenhower C, Huttley GA, Janssen S, Jarmusch AK, Jiang L, Kaehler BD, Kang K Bin, Keefe CR, Keim P, Kelley ST, Knights D, Koester I, Kosciolek T, Kreps J, Langille MGI, Lee J, Ley R, Liu YX, Loftfield E, Lozupone C, Maher M, Marotz C, Martin BD, McDonald D, McIver LJ, Melnik A V., Metcalf JL, Morgan SC, Morton JT, Naimey AT, Navas-Molina JA, Nothias LF, Orchanian SB, Pearson T, Peoples SL, Petras D, Preuss ML, Pruesse E, Rasmussen LB, Rivers A, Robeson MS, Rosenthal P, Segata N, Shaffer M, Shiffer A, Sinha R, Song SJ, Spear JR, Swafford AD, Thompson LR, Torres PJ, Trinh P, Tripathi A, Turnbaugh PJ, Ul-Hasan S, van der Hooft JJJ, Vargas F, Vázquez-Baeza Y, Vogtmann E, von Hippel M, Walters W, Wan Y, Wang M, Warren J, Weber KC, Williamson CHD, Willis AD, Xu ZZ, Zaneveld JR, Zhang Y, Zhu Q, Knight R, Caporaso JG. 2019. Reproducible, interactive, scalable and extensible microbiome data science using QIIME 2. Nat Biotechnol 37:852.

33. Kumpitsch C, Fischmeister FPS, Mahnert A, Lackner S, Wilding M, Sturm C, Springer A, Madl T, Holasek S, Högenauer C, Berg IA, Schoepf V, Moissl-Eichinger C. 2021. Reduced B12 uptake and increased gastrointestinal formate are associated with archaeome-mediated breath methane emission in humans. Microbiome 9:193.

34. Callahan BJ, McMurdie PJ, Rosen MJ, Han AW, Johnson AJA, Holmes SP. 2016. DADA2: High-resolution sample inference from Illumina amplicon data. Nat Methods 13:581–3.

35. Quast C, Pruesse E, Yilmaz P, Gerken J, Schweer T, Yarza P, Peplies J, Glöckner FO. 2013. The SILVA ribosomal RNA gene database project: improved data processing and web-based tools. Nucleic Acids Res 41.

36. Nilsson RH, Larsson KH, Taylor AFS, Bengtsson-Palme J, Jeppesen TS, Schigel D, Kennedy P, Picard K, Glöckner FO, Tedersoo L, Saar I, Kõljalg U, Abarenkov K. 2019. The UNITE database for molecular identification of fungi: handling dark taxa and parallel taxonomic classifications. Nucleic Acids Res 47:D259–D264.

37. Rivers AR, Weber KC, Gardner TG, Liu S, Armstrong SD. 2018. ITSxpress: Software to rapidly trim internally transcribed spacer sequences with quality scores for marker gene analysis. F1000Research 7.

38. Davis NM, Proctor DiM, Holmes SP, Relman DA, Callahan BJ. 2018. Simple statistical identification and removal of contaminant sequences in marker-gene and metagenomics data. Microbiome 6:1–14.

39. Beule L, Karlovsky P. 2020. Improved normalization of species count data in ecology by scaling with ranked subsampling (SRS): application to microbial communities. PeerJ 8.

40. Andrews S. 2010. FASTQC. A quality control tool for high throughput sequence data.

41. Bolger AM, Lohse M, Usadel B. 2014. Trimmomatic: a flexible trimmer for Illumina sequence data. Bioinformatics 30:2114–2120.

42. Langmead B, Salzberg SL. 2012. Fast gapped-read alignment with Bowtie 2. Nat Methods 9:357.

43. Li H, Handsaker B, Wysoker A, Fennell T, Ruan J, Homer N, Marth G, Abecasis G, Durbin R. 2009. The Sequence Alignment/Map format and SAMtools. Bioinformatics 25:2078.

44. Quinlan AR, Hall IM. 2010. BEDTools: a flexible suite of utilities for comparing genomic features. Bioinformatics 26:841–2.

45. Li D, Liu C-M, Luo R, Sadakane K, Lam T-W. 2015. MEGAHIT: an ultra-fast single-node solution for large and complex metagenomics assembly via succinct de Bruijn graph. Bioinformatics 31:1674–6.

46. Wu YW, Tang YH, Tringe SG, Simmons BA, Singer SW. 2014. MaxBin: an automated binning method to recover individual genomes from metagenomes using an expectation-maximization algorithm. Microbiome 2:26.

47. Buchfink B, Xie C, Huson DH. 2014. Fast and sensitive protein alignment using DIAMOND. Nat Methods 2014 121 12:59–60.

48. Beier S, Tappu R, Huson DH. 2017. Functional Analysis in Metagenomics Using MEGAN 6. Funct Metagenomics Tools Appl 65–74.

49. Overbeek R, Olson R, Pusch GD, Olsen GJ, Davis JJ, Disz T, Edwards RA, Gerdes S, Parrello B, Shukla M, Vonstein V, Wattam AR, Xia F, Stevens R. 2014. The SEED and the Rapid Annotation of microbial genomes using Subsystems Technology (RAST). Nucleic Acids Res 42:D206–14.

50. Seemann T. Abricate, Github. https://github.com/tseemann/abricate.

51. Chibani CM, Mahnert A, Borrel G, Almeida A, Werner A, Brugere JF, Gribaldo S, Finn RD, R.A. S, Moissl-Eichinger C. 2021. A catalogue of 1,167 genomes from the human gut archaeome. Nat Microbiol (in Press.

52. Price MN, Dehal PS, Arkin AP. 2010. FastTree 2 – Approximately Maximum-Likelihood Trees for Large Alignments. PLoS One 5:e9490.

53. Vázquez-Baeza Y, Pirrung M, Gonzalez A, Knight R. 2013. EMPeror: a tool for visualizing high-throughput microbial community data. Gigascience 2.

54. Vázquez-Baeza Y, Gonzalez A, Smarr L, McDonald D, Morton JT, Navas-Molina JA, Knight R. 2017. Bringing the Dynamic Microbiome to Life with Animations. Cell Host Microbe 21:7–10.

55. McMurdie PJ, Holmes S. 2013. phyloseq: An R Package for Reproducible Interactive Analysis and Graphics of Microbiome Census Data. PLoS One 8:e61217.

56. Wickham H. 2016. ggplot2: Elegant Graphics for Data Analysis. Springer-Verlag New York. https://ggplot2.tidyverse.org.

57. Mallickid H, Rahnavardid A, Mciverid LJ, Maid S, Zhangid Y, Nguyenid LH, Tickleid TL, Weingart G, Renid B, Schwagerid EH, Chatterjee S, Thompsonid KN, Wilkinsonid JE, Subramanianid A, Lu Y, Waldronid L, Paulsonid JN, Franzosaid EA, Bravo HC, Huttenhowerid C. 2021. Multivariable association discovery in population-scale meta-omics studies. PLOS Comput Biol 17:e1009442.

58. Love MI, Huber W, Anders S. 2014. Moderated estimation of fold change and dispersion for RNA-seq data with DESeq2. Genome Biol 15:1–21.

59. Benjamini Y, Hochberg Y. 1995. Controlling the False Discovery Rate: A Practical and Powerful Approach to Multiple Testing. J R Stat Soc Ser B 57:289–300.

60. DiGiulio DB, Callahan BJ, McMurdie PJ, Costello EK, Lyell DJ, Robaczewska A, Sun CL, Goltsman DSA, Wong RJ, Shawa G, Stevenson DK, Holmes SP, Relman DA. 2015. Temporal and spatial variation of the human microbiota during pregnancy. Proc Natl Acad Sci U S A 112:11060–11065.

61. Livingston EH, Woodward WA, Sarosi GA, Haley RW. 2007. Disconnect between incidence of nonperforated and perforated appendicitis: implications for pathophysiology and management. Ann Surg 245:886–92.

62. Hansson LE, Laurell H, Gunnarsson U. 2008. Impact of time in the development of acute appendicitis. Dig Surg 25:394–399.

63. Walker A, Hatch Q, Drake T, Nelson DW, Fitzpatrick E, Bingham J, Black G, Maykel JA, Steele SR. 2014. Predictors of appendiceal perforation in an equal access system. J Surg Res 190:87–92.

64. Swidsinski A, Loening-Baucke V, Biche-ool S, Guo Y, Dörffel Y, Tertychnyy A, Stonogin S, Sun N-D. 2012. Mucosal invasion by fusobacteria is a common feature of acute appendicitis in Germany, Russia, and China. Saudi J Gastroenterol 18:55.

65. Read E, Curtis MA, Neves JF. 2021. The role of oral bacteria in inflammatory bowel disease. Nat Rev Gastroenterol Hepatol 18:731–742.

66. Brennan CA, Garrett WS. 2019. Fusobacterium nucleatum - symbiont, opportunist and oncobacterium. Nat Rev Microbiol 17:156–166.

67. Warren RL, Freeman DJ, Pleasance S, Watson P, Moore RA, Cochrane K, Allen-Vercoe E, Holt RA. 2013. Co-occurrence of anaerobic bacteria in colorectal carcinomas. Microbiome 1.

68. Bullman S, Pedamallu CS, Sicinska E, Clancy TE, Zhang X, Cai D, Neuberg D, Huang K, Guevara F, Nelson T, Chipashvili O, Hagan T, Walker M, Ramachandran A, Diosdado B, Serna G, Mulet N, Landolfi S, Ramon S, Fasani R, Aguirre AJ, Ng K, Élez E, Ogino S, Tabernero J, Fuchs CS, Hahn WC, Nuciforo P, Meyerson M. 2017. Analysis of Fusobacterium persistence and antibiotic response in colorectal cancer. Science (80-) 358:1443–1448.

69. Finlay BB, Falkow S. 1997. Common themes in microbial pathogenicity revisited. Microbiol Mol Biol Rev 61:136.

70. Rogers AH, Chen J, Zilm PS, Gully NJ. 1998. The behaviour of Fusobacterium nucleatum chemostat-grown in glucose-and amino acid-based chemically defined media. Anaerobe 4:111–6.

71. Kreimeyer A, Perret A, Lechaplais C, Vallenet D, Médigue C, Salanoubat M, Weissenbach J. 2007. Identification of the last unknown genes in the fermentation pathway of lysine. J Biol Chem 282:7191–7197.

72. Di Saverio S, Podda M, De Simone B, Ceresoli M, Augustin G, Gori A, Boermeester M, Sartelli M, Coccolini F, Tarasconi A, De’ Angelis N, Weber DG, Tolonen M, Birindelli A, Biffl W, Moore EE, Kelly M, Soreide K, Kashuk J, Ten Broek R, Gomes CA, Sugrue M, Davies RJ, Damaskos D, Leppäniemi A, Kirkpatrick A, Peitzman AB, Fraga GP, Maier R V., Coimbra R, Chiarugi M, Sganga G, Pisanu A, De’ Angelis GL, Tan E, Van Goor H, Pata F, Di Carlo I, Chiara O, Litvin A, Campanile FC, Sakakushev B, Tomadze G, Demetrashvili Z, Latifi R, Abu-Zidan F, Romeo O, Segovia-Lohse H, Baiocchi G, Costa D, Rizoli S, Balogh ZJ, Bendinelli C, Scalea T, Ivatury R, Velmahos G, Andersson R, Kluger Y, Ansaloni L, Catena F. 2020. Diagnosis and treatment of acute appendicitis: 2020 update of the WSES Jerusalem guidelines. World J Emerg Surg 15:27.

73. Al-Gaithy ZK. 2012. Clinical value of total white blood cells and neutrophil counts in patients with suspected appendicitis: retrospective study. World J Emerg Surg 7:32.

74. Tankel J, Keinan A, Blich O, Koussa M, Helou B, Shay S, Zugayar D, Pikarsky A, Mazeh H, Spira R, Reissman P. 2020. The Decreasing Incidence of Acute Appendicitis During COVID-19: A Retrospective Multi-centre Study. World J Surg 44:2458–2463.

75. Zvizdic Z, Vranic S. 2021. Decreased number of acute appendicitis cases in pediatric population during the COVID-19 pandemic: Any link? J Pediatr Surg. Elsevier https://doi.org/10.1016/j.jpedsurg.2020.08.016.

76. Xiao J, Fiscella KA, Gill SR. 2020. Oral microbiome: possible harbinger for children’s health. Int J Oral Sci 12:1–13.

77. Uchino Y, Goto Y, Konishi Y, Tanabe K, Toda H, Wada M, Kita Y, Beppu M, Mori S, Hijioka H, Otsuka T, Natsugoe S, Hara E, Sugiura T. 2021. Colorectal Cancer Patients Have Four Specific Bacterial Species in Oral and Gut Microbiota in Common—A Metagenomic Comparison with Healthy Subjects. Cancers (Basel) 13:3332.

